# Read Length Dominates Phylogenetic Placement Accuracy of Ancient DNA Reads

**DOI:** 10.1101/2024.06.28.601240

**Authors:** Ben Bettisworth, Nikolaos Psonis, Nikos Poulakakis, Pavlos Pavlidis, Alexandros Stamatakis

## Abstract

A common problem when analyzing ancient DNA (aDNA) data is to identify the species which corresponds to the recovered aDNA sequence(s). The standard approach is to deploy sequence similarity based tools such as BLAST. However, as aDNA reads may frequently either stem from unsampled taxa due to extinction, it is likely that there is no exact match in any database. As a consequence, these tools may not be able to accurately place such reads in a phylogenetic context. Phylogenetic placement is a technique where a read is placed onto a specific branch of a phylogenetic reference tree, which allows for a substantially finer resolution when identifying reads. Prior applications of phylogenetic placement has deployed only on data from extant sources. Therefore, it is unclear how the aDNA damage affects phylogenetic placement’s applicability to aDNA data. To investigate how aDNA damage affects placement accuracy, we re-implemented a statistical model of aDNA damage. We deploy this model, along with a modified version of the existing assessment pipeline PEWO, to 7 empirical datasets with 4 leading tools: APPLES, EPA-ng, pplacer, and RAPPAS. We explore the aDNA damage parameter space via a grid search in order to identify the aDNA damage factors that exhibit the largest impact on placement accuracy. We find that the frequency of DNA backbone nicks (and consequently read length) has the by far largest impact on aDNA read placement accuracy, and that other factors, such as misincorporations, have a negligible effect on overall placement accuracy.

## 1 Introduction

The analysis of DNA obtained from archaeological sites as well as paleontological and sediment samples has changed our understanding of life in the past (Haile et al., 2009; Pedersen et al.,2015; Der Sarkissian et al., 2017). However, the characteristics of ancient DNA (aDNA) samples differ substantially from those of contemporary samples. Due to several degradation processes (including deamination and fragmentation), analyzing aDNA data exhibits more challenges than analyzing DNA obtained from contemporary sources. The DNA repair mechanisms which guarantee that a sample is of high quality become inactive when an organism dies, which allows the DNA to accumulate damage. This accumulated damage often yields poor quality samples.

The damage accumulation rate depends on time, temperature, acidity, and other environmental factors (Sawyer et al., 2012) which complicate the prediction of the *post-mortem* damage level for a given sample. Nonetheless, the damage itself can be described via a straight-forward model, originally proposed by Briggs *et al*. (Briggs et al., 2007). Of the possible damage types DNA accumulates as it ages, the Briggs model of aDNA damage (hereafter, simply the Briggs model) incorporates two major damage types: nicks to the DNA backbone, which shred the DNA into shorter fragments; and point errors caused by cytosine (C) deamination. When cytosine demanites it transforms into uracil (U) and is subsequently read as thymine (T) instead of cytosine (C) during DNA sequencing. This process is commonly known as C to T damage, which we denote as C → T. This damage is observed at the 5’ end of the ancient sequence reads of double-stranded libraries, as well as at both the 5’ and 3’ ends of single-stranded libraries. There exists an analogous G to A damage (which we denote as G → A) that is only observed at the 3’ read ends of double-stranded libraries. This is induced by the genomic library preparation in the wet lab, due to DNA complementarity.

Under the Briggs model, the deamination rate is controlled by the location of the respective base in relation to an overhang. An overhang is a section of the sequence where the complimentary strand has been separated. The bases of the primary strand have therefore been exposed to the environment without the stabilizing presence of the complimentary strand. These overhangs produce so-called single strand regions (in contrast to double strand regions), which exhibit substantially higher rates of deamination due to their unprotected exposure to environmental conditions. For example, Briggs *et. al* found deamination rates in the single strand region to be an order of magnitude higher than in the double strand region. As overhangs necessarily occur at strand ends, the vast majority of C to T damage (and consequently also G to A damage) occurs at the beginning and the end of a read. For an example read of 100 bases, the overhang only needs to be ≈ 3 bases long to exhibit more expected errors than the double stranded region. Many reads are substantially shorter, often only comprising 30 base pairs due to filtering steps after the sequencing process. Hence, due to the short length with respect to the overhang size, most damage occurs at read ends. This induces a characteristic “U” shaped curve of misincorporated point differences as shown in (Briggs et al., 2007; Sawyer et al., 2012).

One major challenge in aDNA data analysis is molecular identification, that is, assigning a DNA read to a taxon. In addition to the processes described above, aDNA samples are also likely to be contaminated by exogenous sources. Determining the organism a read originally belonged to allows researchers to make inferences based on the read context, for example by refining the range, both spatial and temporal, of extinct species (Haile et al., 2009) or uncovering admixture events in archaic hominins, for instance (Slatkin and Racimo, 2016). However, aDNA degradation results in both, small DNA fragments, and erroneous point differences, yielding identification challenging.

Prior data analyses have attempted to assign aDNA reads via tools such as BLAST (Haile et al., 2009) or Kraken (Wood and Salzberg, 2014). These tools match a short (a)DNA query sequence or read against a (possibly large) database of known sequences. However, sequence similarity based methods only match against known sequences, or equivalently, sequences located at the tips of a corresponding phylogenetic tree. However, if the query sequence is sampled from a heretofore unknown, extinct, or hypothetical species, a similarity-based taxonomic assignment will be imprecise. These methods will indicate a single clade, and possibly a confidence associated with the assignment to that clade. Yet, if multiple clades contain a closely matching reference sequence, similarity-based methods will typically only assign the read to a single clade.

An alternative method for taxonomic identifications is to deploy phylogenetic placement, as implemented in EPA-ng, APPLES, `pplacer`, or RAPPAS (Matsen et al., 2010; Barbera et al., 2019; Linard et al., 2019; Balaban et al., 2020). Phylogenetic placement methods place query sequences into a phylogenetic reference tree. Phylogenetic placement is a technique which places unknown, sampled sequences, called query sequences, into a known reference tree inferred from known/named sequences, called reference sequences. By using a phylogeny, placement is able to match query sequences against hypothetical and unsampled species (i.e., an evolutionary history), as well as against known species. Therefore, placement-based taxonomic assignment is often more specific and accurate than a mere assignment to a potentially taxon-rich clade (Berger et al., 2011). This is because phylogenetic placement will assign the query sequence to a specific branch of the phylogenetic reference tree. For a comprehensive review of phylogenetic placement methods that also includes a plethora of references to empirical data analyses on contemporary DNA data please refer to (Czech et al., 2022).

A recent publication raised the question if phylogenetic placement, which was originally devised for analyzing contemporary DNA sequences, can adequately identify aDNA sequences due to their short read length and characteristic damage (Martiniano et al., 2022). However, a systematic investigation of the effects of aDNA damage on phylogenetic placement accuracy has yet to be conducted. In the following, we therefore systematically investigate the impact of aDNA damage on placement accuracy.

To this end, we have modified the existing phylogenetic placement benchmark pipeline PEWO (Linard et al., 2021) by including a step for injecting aDNA damage into the query sequences. Using this modified pipeline we conduct appropriate accuracy tests using the following placement tools: EPA-ng (Barbera et al., 2019), APPLES (Balaban et al., 2020), `pplacer` (Matsen et al., 2010), RAPPAS (Linard et al., 2019).

## 2 Methods

To evaluate the effects of aDNA damage on phylogenetic placement accuracy, we initially reimplemented the popular aDNA damage simulator `Gargammel` (Renaud et al., 2017), as an open source Python program which we call `PyGargammel` (available at https://github.com/computations/pygargammel). This re-implementation is required to better control the particular type and magnitude of aDNA damage (specifically the parameters of the Briggs Model (Briggs et al., 2007) that is implemented in `Gargammel` and `PyGargammel`), as this is difficult or impossible in `Gargammel`. For both simulators, the input is a FASTA file containing one or more sequences. `Gargammel` simulates reads, and therefore outputs the results as FASTQ files, which need to subsequently be assembled *and* aligned (output as BAM) in order to be suitable for use in PEWO. However, as we intend to isolate the effects of aDNA damage exclusively on placement results, our damage simulator implementation allows to skip these potentially confounding steps. To this end, `PyGargammel` outputs its results as sequences in FASTA format, with an option to preserve alignment with the original, undamaged, sequence. We discuss `PyGargammel` in more detail in Section 2.3.

To facilitate running multiple phylogenetic placement programs on alignments with simulated damage from PyGargammel, we used the placement accuracy assessment pipeline PEWO (Linard et al., 2021). However, executing PEWO with PyGargammel, required several modifications to PEWO which we detail in Section 2.4.

Finally, several damage simulation parameters needed to be chosen for this application. Instead of selecting single parameters or sampling the parameter space uniformly, we chose to perform a grid search over possible damage parameter configurations. These choices are detailed in Section 2.5.

### 2.1 Data

We investigated the effects of aDNA damage on placement accuracy using seven empirical datasets. The dataset properties, such as taxon number and site count, are provided in Table 1. We sought to include a wide range of datasets, including standard phylogenetic placement applications for environmental DNA (Srinivasan et al., 2012; Sunagawa et al., 2015; Mahé et al., 2017), but also datasets from prior aDNA studies. These datasets comprise for the most part larger animals, such as the Aurochs (Gurke et al., 2021), Land Snails (Psonis et al., 2022a), Hippos (Psonis et al., 2022b), and Elephants (Psonis et al., 2020). These datasets therefore represent a broad spectrum of aDNA identification tasks. We chose to use seven datasets in order to focus on parameter exploration, rather than expending a large number of resources on a shallow exploration of many datasets. Nonetheless, we have sought to include diverse enough datasets so that many typical aDNA data applications are well represented by our results.

**Table 1:**
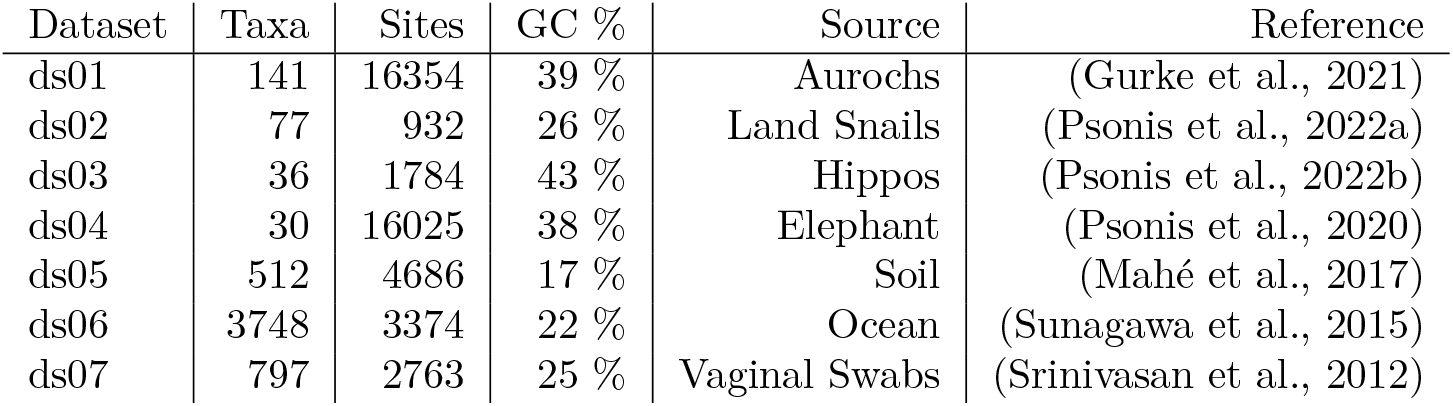
Table of empirical datasets used for experiments. Source is a short description of either the environment the sample was collected from, or the taxonomic clade the samples represent. GC % is the average GC content for all sequences in the alignment.

In addition to 7 empirical datasets, we also investigated the impact of GC content on placement accuracy in the presence of aDNA damage using 9 simulated datasets. The description of how these datasets were generated, as well as the results from these datasets are described in the Supplemental Material.

### 2.2 The PEWO Pipeline

Instead of re-implementing an evaluation pipeline for analyzing phylogenetic placement accuracy, we appropriately modified the existing PEWO pipeline (Linard et al., 2021) to also support damage simulation. Therefore, we enhanced PEWO by directly integrating simulation of damaged sequences via a damage simulator we call PyGargammel. The details of PyGargammel and how it simulates damage are found in Section 2.3.

Given a multiple sequence alignment (MSA) and an associated phylogeny, PEWO performs analysis by

**Pruning** Pruning the taxa contained in a randomly selected subtree, as well as removing and fragmenting the sequences contained in this subtree from the MSA,

**Alignment** Realigning the pruned and shortened sequences all at once with HMMER (Eddy, 2011),

**Placement** Running various placement programs to place the pruned sequences back on the reference tree,

**Assessment** and Computing and summarizing the placement error with regards to the ground truth placement.

PEWO is designed to investigate the accuracy of phlyogentic placement tools when given somewhat realistic data. To this end, it will fragment the pruned portion of the input MSA into multiple parts. However, the behavior of aDNA fragmentation is very different from how sequence fragmentation is implemented in PEWO. Therefore, we implement our own fragmenting method (detailed in Section 2.3), and disable the sequence fragmentation in PEWO.

### 2.3 Simulating Ancient DNA Damage

To simulate the characteristic *post-mortem* damage that DNA undergoes as it ages, we implemented the Briggs Model of DNA damage (Briggs et al., 2007) in a tool we call PyGargammel. The primary use case of PyGargammel is to take a FASTA file with sequences and insert aDNA typical damage. This model has four parameters which control the effects and magnitude of aDNA-like sequence damage. Using the same notation as Briggs et al. (2007), the parameters are:

- The probability of nicks 0 ≤ *λ* ≤ 1,
- the parameter for the geometric distribution from which the overhang length is drawn 0 < *v* ≤ 1,
- the probability of deamination in single stranded regions 0 ≤ *δ*_*ss*_ ≤ 1,
- and the probability of deamination in double stranded regions 0 ≤ *δ*_*ds*_ ≤ 1.

The first mechanism in the model determines the probability of breaking (nicking) *λ*, for each base. These reads then have an overhang with a length that is drawn from geometric distribution with success probability *ν*. Which side of the reads (5’ or 3’) the overhang is located on depends on numerous factors which occur during sequencing. However, for the purpose of damage simulation, it is essential that the overhang is present only on the 5’ or the 3’ side of the read. Therefore, when simulating the overhang, we randomly generate an overhang on the 5’ side for 50% of reads, and generate an overhang on the 3’ side for the remaining 50% of the reads. Then, deamination damage is applied with a probability that depends on whether the base is in a single or double stranded region. If the base is in the overhang region, then the single stranded deamination rate *δ*_*ss*_ is used to determine the deamination probability. If the base is not located in an overhang region, then the double stranded deamination rate *δ*_*ds*_ applies.

Note that there are two distinct types of possible deamination damage, C → T and G → A. C → T damage is the direct result of a cytosine being deaminated (this is to say, losing a part of its chemical structure) which results in the base being misincorporated as a T. However, G → A damage is *not* a direct result of deamination. Instead, it is a consequence of the genomic library preparation for the sequencing process. More specifically, when the sample is prepared for sequencing, overhangs are repaired, which transforms the G → A damage into C → T damage on the complimentary strand. In this case, the overhang is repaired with an A instead of a G, as G is the complimentary base to a deaminated C. If the sequence is read from left to right, then *only* C → T damage can occur in the read if the overhang is on the left side, and *only* G → A damage occurs if the overhang is on the right side.

PyGargammel relies on this model and implements it as follows: It starts by reading a FASTA file and iterates over each sequence in the file by performing the following steps:

1. Iterate through bases and mark nicks with probability *λ*.
2. Break the sequence into reads at the nick points.
3. Compute overhang lengths on both ends with distribution Geom(*p* = *ν*).
4. Perform deaminations (C → T and G → A damage)
  - If the read has an overhang (with a possible length of zero) on the 5’ end, and the base is in the overhang region, apply C → T damage with probability *δ*_*ss*_. If the base is not in the overhang region, apply C → T damage with probability *δ*_*ds*_.
  - Likewise, if the read has an overhang (with a possible length of zero) on the 3’ end, and the base is in the overhang region, apply G → A damage with probability *δ*_*ss*_. If the base is not in the overhang region, apply damage with probability *δ*_*ds*_.
5. (Optional) Position the read such that it is aligned with its original position in the original sequence.
6. (Optional) Filter sequences that are too short if a minimum read length was supplied.
7. (Optional) If a minimum read count threshold was specified, repeat the process from step 1 accumulating additional reads until the threshold has been achieved.
8. (Optional) If a maximum number of reads was specified, and too many reads were generated, downsample uniformly at random to the provided threshold.

It is important to discuss how PyGargammel handles ambiguous characters and missing data, as empirical MSAs often have ambiguous characters which can affect the results of downstream analysis. As PyGargammel is designed to also operate on already aligned data, there might be gaps present in the sequences to be damaged. The resulting reads need to be still in alignment after processing, so gaps and ambiguous characters are retained as the respective sequence is being processed. This means that gap characters will appear in the resulting damaged reads, and gaps will also affect the damage distribution, as gap characters cannot undergo either C → T or G → A damage. However, PyGargammel will completely remove reads entirely consisting of either fully ambiguous or gap characters. In addition, when a minimum read length is specified, only non-gap sites are counted to determine read length.

Finally, if a reference alignment is provided, PyGargammel will reject reads which have no non-gap characters in common, a technique known as premasking. We believe this choice is justified, as such a reads would be difficult to align, and therefore difficult to classify by any method.

### 2.4 Modifications to the PEWO Pipeline

We modified PEWO by inserting PyGargammel both before and after the alignment step. Injecting aDNA sequence damage before alignment is more realistic as aligning reads is a necessary step prior to placement for empirical sequences. However, we also inject aDNA damage *after* the alignment stage, so as to isolate the effects of aDNA damage on placement accuracy, without any confounding alignment error. Unless otherwise stated, we report placement accuracy for damage injection *after* alignment. We did conduct some exploratory experiments with damage inserted before alignment, however a thorough investigation of the joint alignment and placement error is beyond the scope of this paper. Specifically, we wish to investigate the case when alignment error is negligible to determine the baseline effectiveness of placement algorithms.

Therefore, the operation order for our analysis pipeline is (unless otherwise stated).

**Pruning** Pruning the taxa associated with a random subtree, as well as removing the sequences from the MSA,

**Alignment** Realigning the pruned sequences all together with respect to the remaining reference MSA,

**Damage** Fragmenting and deaminating the aligned sequences,

**Placement** Running various placement programs on the pruned sequences,

**Assessment** and Computing and summarizing the placement error.

We have also prepared a graphical depiction of these steps which is presented in Figure 1. The full pipeline can be found on GitHub at https://github.com/computations/PEWO

**Figure 1:**
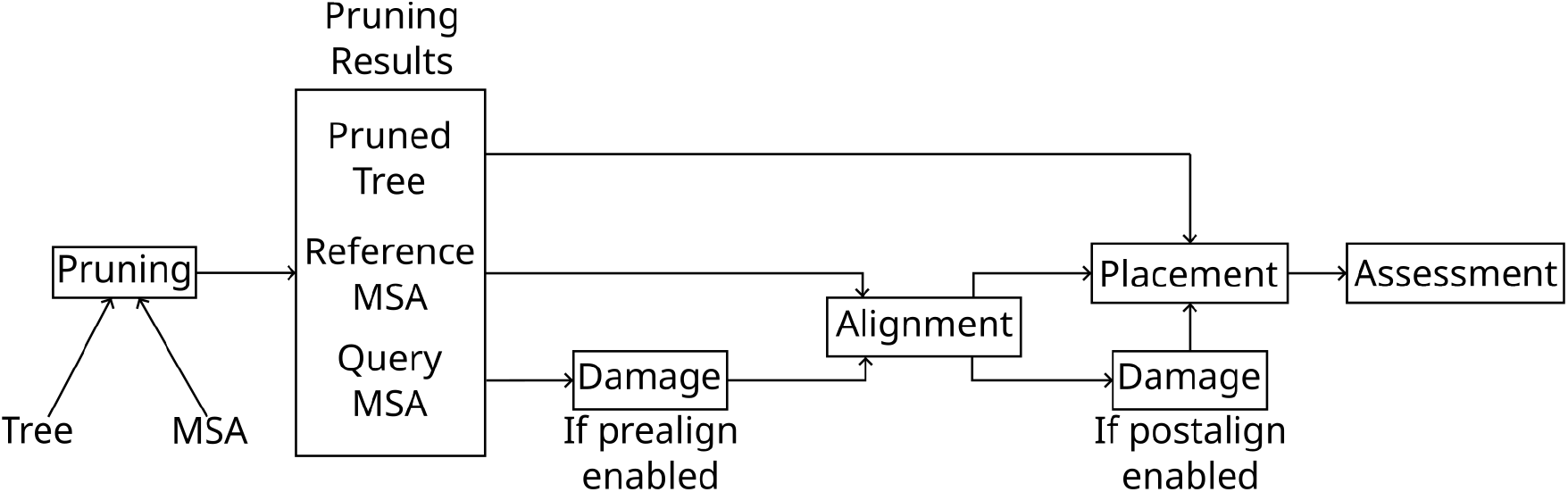
Diagram of the modified PEWO pipeline. The steps “Pruning”, “Alignment”, “Damage”, “Placement”, and “Assessment” are described in Section 2.3.

### 2.5 Pipeline Parameters

In order to fully investigate aDNA damage impact on phylogenetic placement accuracy, we perform a grid search over the damage simulation parameter configurations. We based the endpoints of this grid search for each parameter on the estimated values from Briggs et al. (2007). The specific values that were tested are provided in Table 2.5. If we increase the number of investigated values per parameter equally, then the number of evaluated points grows with*≀*(*n*^4^). This means that the number of points to be evaluated quickly becomes too large to be computed in reasonable time. We can be more efficient if we investigate the behavior and impact of a single parameter at a time while keeping the remaining parameters fixed. An initial parameter space exploration with a single dataset showed that the nick frequency is the most important factor (w.r.t. its impact on placement accuracy). Hence, we thoroughly investigate the effects of the nick frequency parameter, only. To this end, we sample *λ* values ranging from0.0 to 0.03. The value of 0.03 slightly exceeds the value estimated in Briggs et al. (2007), which was found to be *λ* = 0.024 for Neanderthal samples. Overall, we assessed nine *λ* values, with a higher density near 0.0, as that is where placement accuracy is most sensitive to small changes in *λ*. In Table 3 we show both the theoretical and the actual median read length for each value of *λ* in order to better illustrate the impact of changes in *λ* on read length. Noticeably, the largest reductions in both, estimated, and actual median read length stem from values near 0.0. The values differ between the estimated values and the actual values due to finite data, as the maximum length of any read is limited by the original sequence length. Furthermore, we also exclude reads with less than 15 base pairs, which further skews the actual values.

**Table 2:**
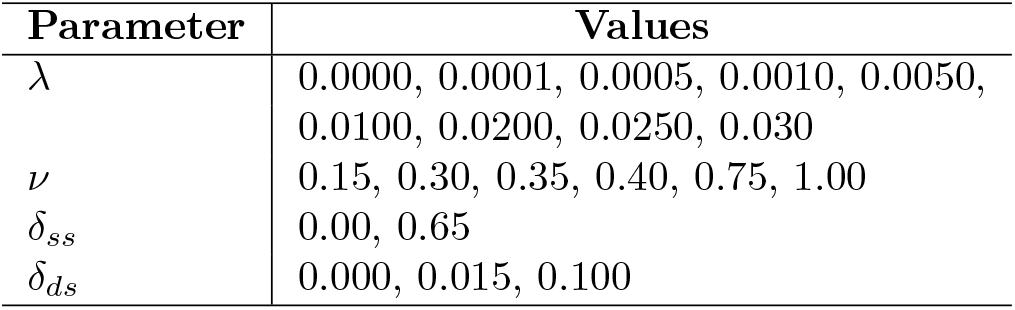
Table containing the parameter values used in our experiments. End points for each parameter are based on the values estimated in Briggs et al. (2007).

**Table 3:** Tool option and heuristics used for all datasets and parameter values.

| Tool | Options and Heuristics |
| --- | --- |
| RAPPAS | `k = 7, omega = 2.0, reduction = 0.99` |
| EPA-NG | `h1, g = 0.99999` |
| pplacer | `max-strikes = 6, strike-box = 3, max-pitches = 40` |
| APPLES | `OLS, MLSE` |

Beyond *λ*, we experimentally determined that the overhang parameter *ν* is the second most important parameter, again with respect to its impact on placement accuracy. As before, we base the endpoints of grid values on those listed in Briggs et al. (2007). Since overhangs are modeled as iteratively “growing” with probability *ν* of the process terminating at the current overhang length, the parameter value associated with no damage is 1.0. This is when the process terminates immediately, resulting in a zero length overhang. Correspondingly, the overhang lengths increase as the parameter value decreases. We tested six values with the most extreme damage being 0.15, which is slightly more extreme than the value estimated in Briggs et al. (2007).

The deamination rates *δ*_*ss*_ and *δ*_*ds*_ were both found to have a minor damage effect during initial experiments, therefore we only consider two values for *δ*_*ss*_ and three values for *δ*_*ds*_ in our grid. For *δ*_*ss*_ the upper endpoint follows, again, Briggs et al. (2007). On the other hand, the upper *δ*_*ds*_ value is a very extreme value of 0.1, which is far greater than the value found in Briggs et al. (2007). We included this extreme value because we intend to investigate the effects of high levels of C → T and G → A damage on placement accuracy, as this is the unique feature that yields aDNA damage being different from “just” very short reads. In particular, we explore placement accuracy for damage rates that are substantially higher than expected in order to investigate the maximal effect of misincorperation on placement accuracy.

Finally, we implemented two additional constraints on the simulated reads. The first is thata minimum of 10 reads per selected sequence from the reference MSA is generated. This ensures that we have a sufficient number of reads to compute proper summary statistics. The second is a minimum read length of 15 base pairs. This mimics representative analyses of aDNA samples, where reads under a certain length (typically 30 base pairs) are excluded in a prefiltering step.

### 2.6 Analysis

We ran PEWO with the following placement tools: RAPPAS(Linard et al., 2019), EPA-`ng`(Barbera et al., 2019), `pplacer`(Matsen et al., 2010), and APPLES(Balaban et al., 2020). We do not explore different placement program options (e.g., various heuristics) because initial experiments did not exhibit substantial placement accuracy differences as a function of these options. Therefore, in order to also economize on computational resources, we elected to use one set of placement program options for all runs. These options are described in Table 3.

Experiments were performed on a server with 40 x86 CPU^1^ cores and 754 GiB of memory. The number of independent subtree prunings and accuracy evaluations (denoted as a “pruning” in PEWO) was set to 10 for all datasets. This is to say, that 10 separate experiments were conducted for each set of parameters on each reference MSA and the results were aggregated. PEWO was run with 40 cores.

To better visualize our experimental results, we fit a linear model augmented with cubic splines for each parameter against the expected node distance (eND). The expected node distance is computed as LWR(*p*)×*d*(*p*) where, LWR(*p*) is the likelihood weight ratio of the placement, and *d*(*p*) is the node distance between the estimated placement and the true placement. We use eND as it better accounts for uncertainty in the specific placement location. We added splines to the model in order to capture the decreasing marginal impact of the damage model parameters. The regression was performed using R, with the command geom_smooth(method=“lm”, formula = y ∼ splines::ns(x, *k*)) where *k* is the number of splines used for the regression on the specific parameter. We used *k* := 3 for the regression on *λ* and *ν, k* := 2 for the regression on *δ*_*ds*_, and *k* := 1 for the regression on *δ*_*ss*_. We vary the number of splines because the number of parameter values sampled varies between parameters.

## 3 Results

The regressions describing placement accuracy as a function of different simulation parameter values are displayed in Figure 4. Additionally, the *R*^2^ for each regression is shown in Table 4. We found that the primary factor affecting placement accuracy is the nick frequency parameter *λ*. As seen in Figure 4 and in Table 4, the only parameter which consistently impacts placement accuracy (as measured by expected node distance) is *λ*. For all regressions, the variance explained due to variations in *λ* exceeded that of all other parameters by at least one order of magnitude.

**Table 4:** Table of *R*^2^ values for the regressions in Figure 4. Additionally shown is the *R*^2^ for the regression with all datasets included. Values were computed with lm(e_nd ∼ splines::ns(*p, k*)) where *p* is one of the parameters, and *k* is the number of splines used for that parameter’s model. We used *k* := 3 for *λ* and *v, k* := 2 for *δ*_*ds*_, and *k* := 1 for *δ*_*ss*_.

| Dataset | $R^2$ | | | |
| --- | --- | --- | --- | --- |
| | $\lambda$ | $\nu$ | $\delta_{ss}$ | $\delta_{ds}$ |
| ds01 | 0.0805 | $1.388 \times 10^{-4}$ | $9.199 \times 10^{-6}$ | $2.125 \times 10^{-3}$ |
| ds02 | 0.1415 | $1.687 \times 10^{-5}$ | $1.896 \times 10^{-5}$ | $3.590 \times 10^{-4}$ |
| ds03 | 0.1925 | $1.557 \times 10^{-4}$ | $2.973 \times 10^{-5}$ | $1.358 \times 10^{-3}$ |
| ds04 | 0.1996 | $4.426 \times 10^{-5}$ | $1.896 \times 10^{-5}$ | $1.020 \times 10^{-5}$ |
| ds05 | 0.1498 | $1.099 \times 10^{-5}$ | $3.374 \times 10^{-6}$ | $2.613 \times 10^{-3}$ |
| ds06 | 0.2006 | $1.101 \times 10^{-5}$ | $3.147 \times 10^{-5}$ | $1.648 \times 10^{-4}$ |
| ds07 | 0.1007 | $5.732 \times 10^{-5}$ | $1.132 \times 10^{-4}$ | $5.227 \times 10^{-3}$ |
| All | 0.0610 | $5.408 \times 10^{-5}$ | $2.230 \times 10^{-7}$ | $2.537 \times 10^{-4}$ |

**Table 5:** Table of estimated and actual read lengths based on the Briggs model. The median is computed for each dataset. Discrepancy between the estimated median and the actual median is due to: the finite length of the data; and the minimum read length specified as part of the pipeline.

| $\lambda$ | Median Read Length | |
| --- | --- | --- |
|  | Estimated | Actual |
| 0.0000 | ∞ | 3409 |
| 0.0001 | 6931 | 3388 |
| 0.0005 | 1385 | 1309 |
| 0.001 | 692 | 736 |
| 0.005 | 138 | 165 |
| 0.01 | 68 | 91 |
| 0.02 | 34 | 53 |
| 0.025 | 27 | 45 |
| 0.03 | 22 | 40 |

The parameter *δ*_*ds*_ had the second most substantial impact on placement accuracy. Nonetheless, it is almost negligible compared to the impact of *λ*. For all datasets, the *R*^2^ for the regressionwith *δ*_*ds*_ was at least 1 order of magnitude smaller than the *R*^2^ for the regression with *λ*.

The remaining parameters had a negligible effect on placement accuracy. The *R*^2^ for all models was at least two orders of magnitude smaller for all datasets. To further investigate this, we computed the error rate for generated reads. In Figure 5 we show the error rate as a histogram for datasets with *λ* = 0.025,*ν* = 0.15, *δ*_*ss*_ = 0.65 and *δ*_*ds*_ = 0.015. Most reads exhibit a relatively small number of errors, with a median error rate of 4%.

To better illustrate the effect of a high nick rate on placement accuracy we plotted a histogram of eND normalized by tip count in Figures 2 and 3.

**Figure 2:**
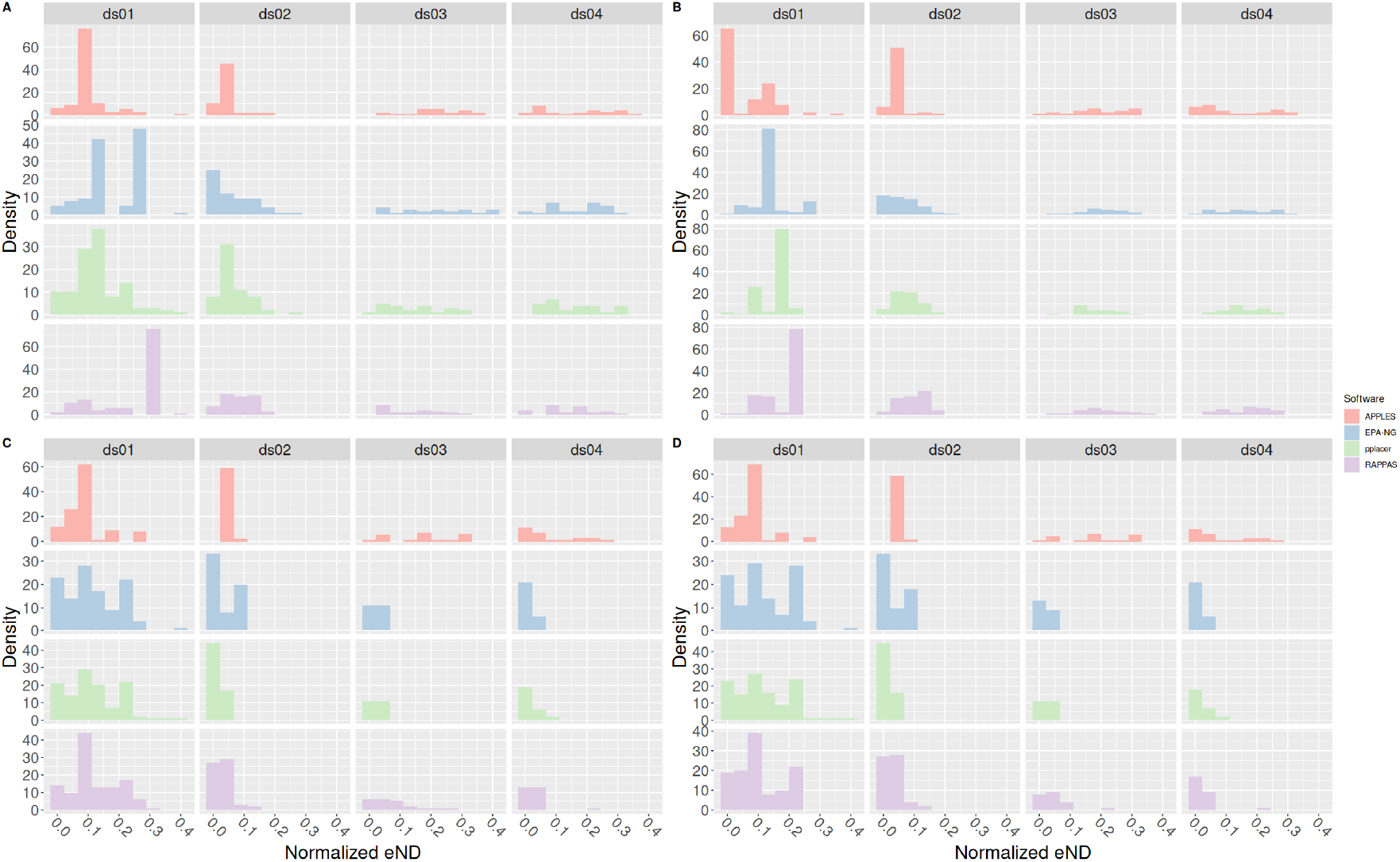
Histogram of normalized effective node distances for datasets ds01, ds02, ds03, and ds04. In addition, each subfigure has been faceted by the respective placement software being used. Top plots (A and B) have nicks enabled (*λ* = 0.025), and bottom plots (C and D) have nicks disabled (*λ* = 0.0). Plots on the left (A and C) have other forms of damage enabled (*ν* = 0.15, *δ*_*ss*_ = 0.65, *δ*_*ds*_ = 0.015). Plots on the right (B and D) have other forms of damage disabled (*ν* = 1.0, *δ*_*ss*_ = *δ*_*ds*_ = 0.0). All plots share the same scale for the x-axis.

**Figure 3:**
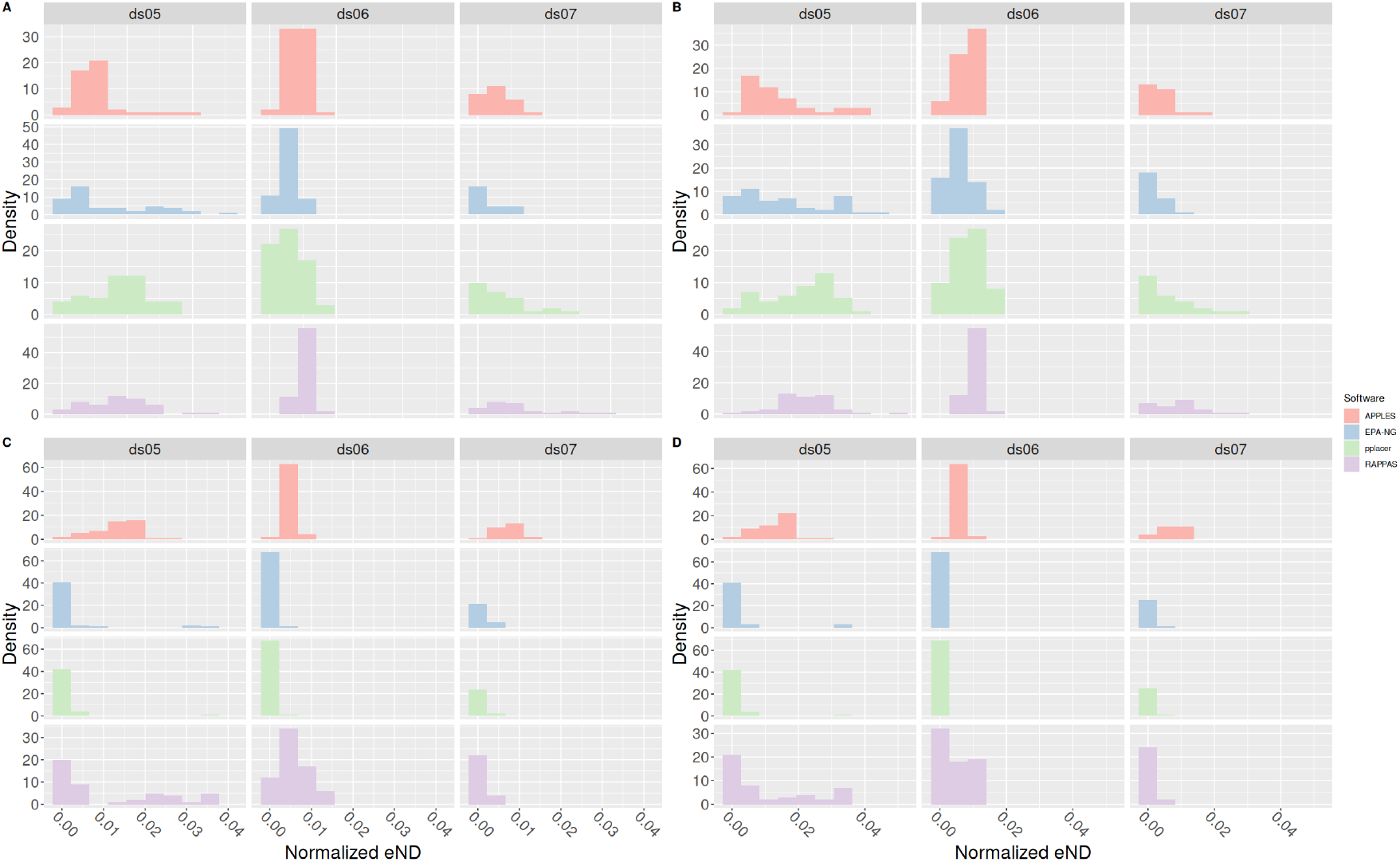
Histogram of normalized effective node distances for datasets ds05, ds06, and ds07. In addition, each subfigure has been faceted by the placement software used. Top plots (A and B) have nicks enabled (*λ* = 0.025), and bottom plots (C and D) have nicks disabled (*λ* = 0.0). Plots on the left (A and C) have other forms of damage enabled (*ν* = 0.15, *δ*_*ss*_ = 0.65, *δ*_*ds*_ = 0.015). Plots on the right (B and D) have other forms of damage disabled (*ν* = 1.0, *δ*_*ss*_ = *δ*_*ds*_ = 0.0). All plots share the same scale for the x-axis.

## 4 Discussion

Based on our analysis, the primary factor influencing aDNA placement accuracy is the nick frequency, or equivalently the read length. In Figures 2 and 3, for example, we can see a clear shift to the left between the top and bottom (that is, the plots with *λ* = 0.025 vs the plots with *λ* = 0.0). This shift to the left indicates an average improvement in placement accuracy. In contrast, we do not observe a similar shift between the left (other damage types enabled) and right (other damage types disabled) plots. This indicates, again, that the primary driver of placement accuracy is nick frequency, and that the effects of deamination errors are minor, confirming our repeated empirical observations that likelihood-based models as implemented in EPA-`ng` are generally robust with respect to noise and sequencing error.

One way to ameliorate the placement error from aDNA damage is to incorporate the aging process as an additional model process. Currently, none of the placement tools tested here do this. However, the largest contributor to placement error when analysing aDNA data is the amount of information available in the sequence to be placed. More simply, read length is the primary driver of error for aDNA reads. When reads become shorter, the probability of correctly placing the reads decreases (e.g. Fig 4). The reason for this becomes clear if we examine the distribution of read lengths as shown in Table 3. Both the estimated and actual median read lengths rapidly decrease as *λ* increases. This reflects the increase in placement error shown in Figure 4**A** with increasing nick frequency.

**Figure 4:**
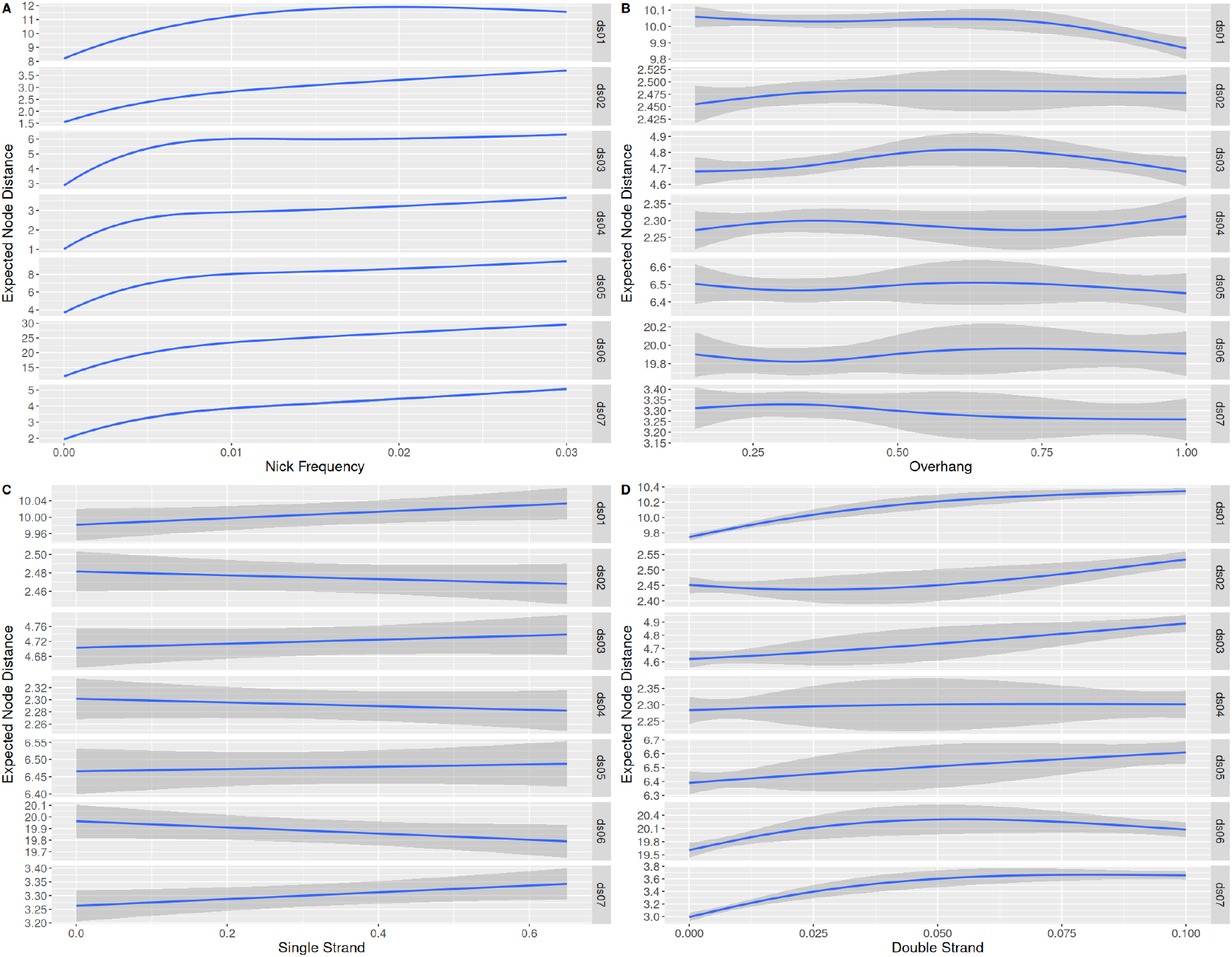
Plot of regressions against parameters for each dataset. Please note that the y scale varies for each dataset. Each regression is against the expected node distance (eND). The subfigures **A, B, C**, and **D** plot *λ, v, δ*_*ss*_m and *δ*_*ds*_ against eND, respectively. Plot generated using ggplot2 with `geom_smooth(method=“lm”, sd=TRUE, formula = y ∼ splines::ns(x,` *k*)), where *k* varies with the dataset. *k* = 3 for plots **A** and **B**, *k* = 2 for plot **D**, and *k* = 1 for plot **C**.

In contrast, the total amount of deamination events (that is, C → T and G → A damage)is not as important for placement accuracy. As we have shown in Figure 5, the rate of site errors is relatively low compared to the length of the read, even for higher damage rates. This is expected, as only two bases (C and G) are available to being misincorporated. On a typical “random” sequence, only 50% of sites will be eligible to be damaged. Furthermore, in practice GC content is often below Of course, the so-called GC-content varies among species and genomic regions. Hence we would expect that sequences will be more difficult to place correctly as a function of their GC-content. However, we investigated the effect of increasing amounts of GC-content on placement accuracy in the presence of aDNA damage using simulated data, and found no major differences between high and and low GC-content datasets. Details of this analysis are described in the Supplemental Material.

**Figure 5:**
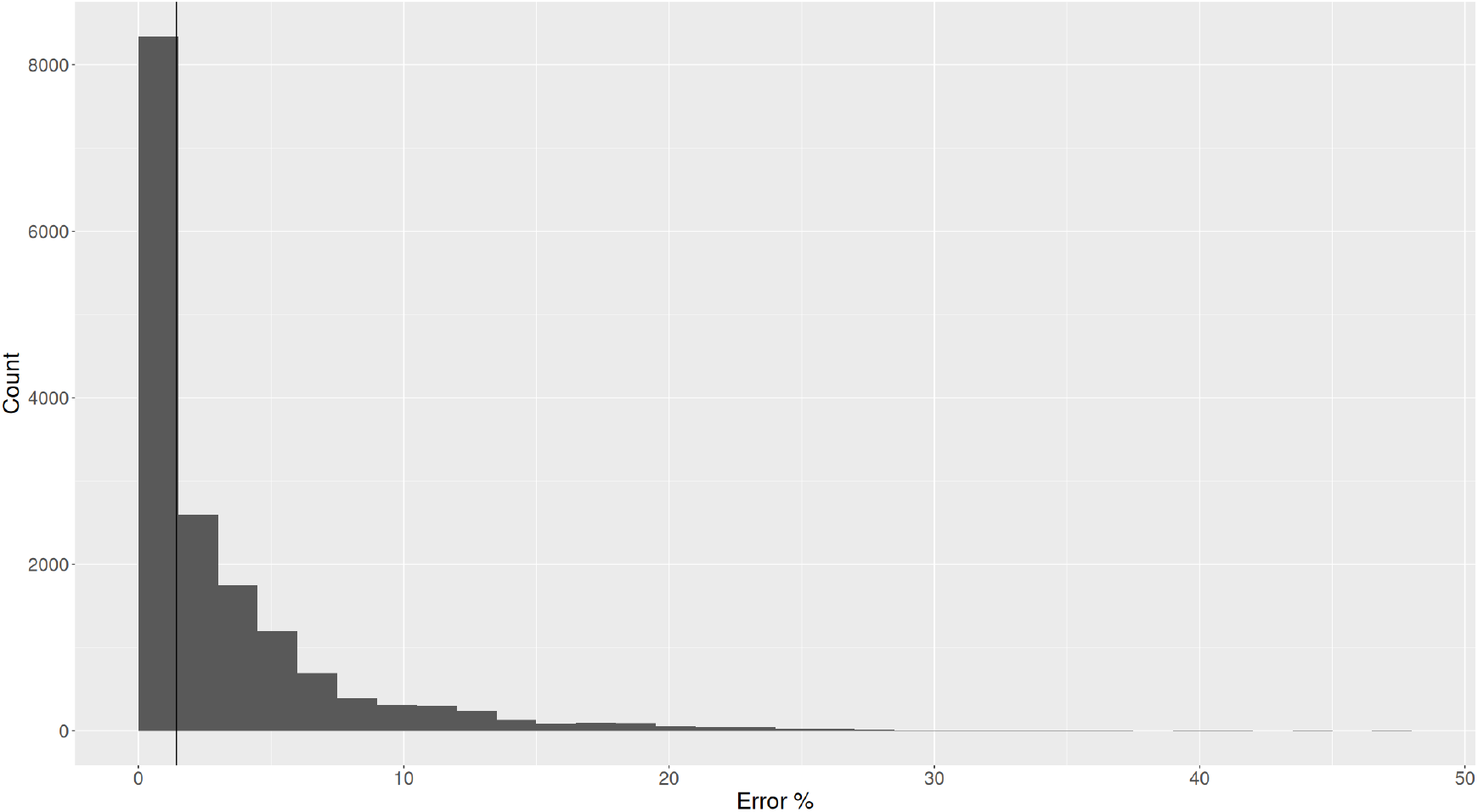
Histogram of percent of error sites per read for reads generated under the model parameters *λ* = 0.025, *v* = 0.15, *δ*_*ss*_ = 0.65 and *δ*_*ds*_ = 0.015. An error site is a site which has either experienced C → T or G → A damage. A vertical line is drawn at the median, where 1.4% of sites are errors. The max percent error for all reads is 47%

While we based our investigation around parameters found in Briggs et al. (2007), which is based on reads obtained from Neanderthal samples, the parameter values tested also cover other data sources, such as molluscs (Pedersen et al., 2015). Additionally, further investigation into some of the parameters is unlikely to clarify the issue any further, especially for the nick frequency *λ*. At the maximum value of *λ* in this study, many of the reads generated were filtered by the already permissive (w.r.t. to current practice in aDNA studies) minimum read length cutoff at 15 sites. Further increases in *λ* would only marginally decrease the realized read lengths, as most reads would be filtered by the minimum read length.

All of this together indicates that phylogenetic placement is an appropriate tool for analysing aDNA sequence. While the frequency of nicks strongly affects the probability of accurately placing reads, this is predominantly due to length of the reads to be placed, and not to other damage types. Importantly, *all* methods to identify aDNA reads will experience difficulties with extremely short reads, as the amount of information contained within a read is directly proportional to its length. Furthermore, placement seems to be robust to other forms of aDNA damage, even when reads are short (see Figures 2 and 3.)

There are some limitations to this work. First, we did not examine the case where C → T damage occurs in both overhangs instead of G → A in the second overhang, as is the case in single stranded libraries. However, we do not expect this to have any effect on the results presented here, as it is likely the number of misincorporations that matters, and not the type. Additionally, we did not thoroughly examine the effects of alignment on placement accuracy, even though we might expect alignment to strongly affect results. In particular, we used the multiple sequence aligner HMMER (Eddy, 2011), which aligns all sequences simultaneously. Another method, which we did not explore, is to conduct the simpler step of single sequence alignment, where only one read is aligned to the reference MSA at a time. In contrast to multiple sequence alignment, single sequence alignment does not change the reference alignment, and so it might be more robust to mis-aligning common errors like C → T or G → A damage.

Finally, our work indicates that a dedicated aDNA damage-aware phylogenetic placement model is unlikely to substantially improve results.

## Data Availability

The modified PEWO pipeline is availible at github.com/computations/PEWO. Datasets, result files, plotting scripts, and utility scripts used can be found

## Acknowledgment

This work was funded by the European Union (EU) under Grant Agreement No 101087081 (Comp-Bio- div-GR).

## Footnotes

\# specifically a dual socket server with 2 Intel Xeon Gold 6148

