## Supplemental Text for "Read Length Dominates Phylogenetic Placement Accuracy of Ancient DNA Reads"

### Supplemental Material for "Read Length Dominates Phylogenetic Placement Accuracy of Ancient DNA Reads"

June 28, 2024

#### 1 Investigating the effects of GC content on placement error in the presence of aDNA damage

In order to be sure that our results are not driven by our utilization of GC-poor datasets, we simulated datasets with varying levels of GC content. To generate the sequences we generated random trees via a custom tool which uses a random coalescent process (similar to `rcoal`). Using this random tree, we used AliSim (Ly-Trong et al.) to generate a simulated sequence using the following command.

```
iqtree2
    --alisim <PREFIX>
    -t <RANDOM TREE>
    --out-format fasta
    --length 5000
    -m GTR+G+I+F
      {(.5 - .05*i)/(.05 * i)/(.05 * i)/(.5 - .05 * i)}
```

This simulates a 5000 b.p. sequence under the GTR model (with equal rates), 4 gamma rate categories and invariant sites. Initial frequencies are based on the dataset name, where `i` is the value of `sds0i`. For example, `sds03` has initial frequncies of `{(0.5 - 0.05 * 3) / (0.05 * 3) / (0.05 * 3) / (0.5 - 0.05 * 3)}` which is equal to `{0.35 / 0.15 / 0.15 / 0.35}`. Each data dataset was run using the modified PEWO pipeline as described in Section 2 of the main text, except that the tool `pplacer` was skipped due to technical difficulties. Results from these runs are summarized in Figure 1. The regressions parameters are the same as in Figure 4 of the main text.

As it can be seen Figure 1, the GC content of the data does not impact the results. The major conclusion of the main paper, that read length is the single most important factor impacting phylogenetic placement accuracy, is unaffected by the GC content of the data.

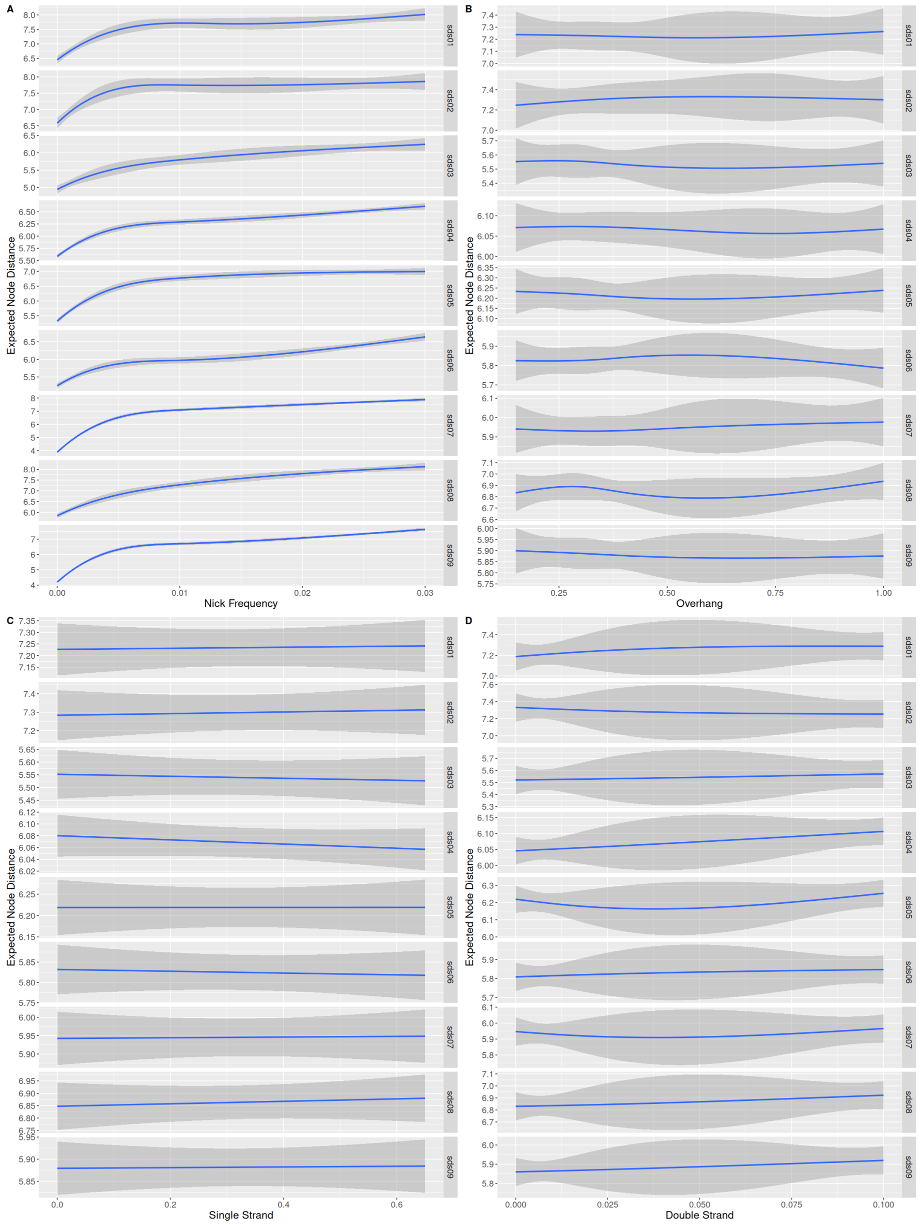


Supplementary Figure 1: Regressions for model parameters using simulated data with varying levels of GC content. GC content is minimized in sds01 (GC% of 10%), and increases by steps of 10% until it is maximized in sds09 (GC% of 90%). Sequences were simulated using AliSim (Ly-Trong et al.).

| Dataset | Initial Frequencies | Realized GC% |
|---|---|---|
| sds01 | {.45 / .05 / .05 / .45 } | 10.1% |
| sds02 | {.40 / .10 / .10 / .40 } | 19.9% |
| sds03 | {.35 / .15 / .15 / .35 } | 29.9% |
| sds04 | {.30 / .20 / .20 / .30 } | 39.9% |
| sds05 | {.25 / .25 / .25 / .25 } | 50.0% |
| sds06 | {.20 / .30 / .30 / .20 } | 60.0% |
| sds07 | {.15 / .35 / .35 / .15 } | 69.9% |
| sds08 | {.10 / .20 / .20 / .10 } | 80.0% |
| sds09 | {.05 / .45 / .45 / .05 } | 90.0% |

Table 1: Datasets, the initial frequency parameters used for simulation, and the realized GC% of each dataset.

### References


Nhan Ly-Trong, Suha Naser-Khdour, Robert Lanfear, and Bui Quang Minh. Alisim: A fast and versatile phylogenetic sequence simulator for the genomic era. *Molecular Biology and Evolution*, 39. doi: 10.1093/molbev/msac092. URL `https://dx.doi.org/10.1093/molbev/msac092`.
